# Decoding the elite soccer player’s psychological profile

**DOI:** 10.1101/2024.08.25.609552

**Authors:** Leonardo Bonetti, Torbjōrn Vestberg, Reza Jafari, Debora Seghezzi, Martin Ingvar, Morten L. Kringelbach, Alberto Goncalves, Predrag Petrovic

**Affiliations:** Center for Music in the Brain, Department of Clinical Medicine, Aarhus University & The Royal Academy of Music, Aarhus, Aalborg, Denmark; Centre for Eudaimonia and Human Flourishing, Linacre College, University of Oxford, Oxford, United Kingdom; Department of Psychiatry, University of Oxford, Oxford, United Kingdom; Department of Clinical Neuroscience, Karolinska Institutet, Stockholm, Sweden; Department of Psychology, University of Bologna; Psychological Sciences, University of Gloucestershire, School of Natural, Social and Sport Sciences, QU214, Francis Close Hall Campus, Swindon Road, Gloucestershire, Cheltenham GL53 7JX, UK

**Keywords:** Cognitive abilities, Personality, Soccer, Artificial neural networks

## Abstract

Soccer is arguably the most widely followed sport worldwide, and many dream of becoming soccer players. However, only a few manage to achieve this dream, which has cast a significant spotlight on elite soccer players who possess exceptional skills to rise above the rest. Originally, such attention was focused on their great physical abilities. However, recently, it a new perspective has emerged, suggesting that being an elite soccer player require a deep understanding of the game, rapid information processing and decision-making. This growing attention has led to several studies suggesting higher executive functions in soccer players compared to the general population. Unfortunately, these studies often had small and non-elite samples, focusing mainly on executive functions alone without employing advanced machine learning techniques. In this study, we used artificial neural networks to comprehensively investigate the personality traits and cognitive abilities of a sample of 328 participants, including 204 elite soccer players from the top teams in Brazil and Sweden. Our findings indicate that elite soccer players demonstrate heightened planning and memory capacities, enhanced executive functions, especially cognitive flexibility, elevated levels of conscientiousness, extraversion, and openness to experience, coupled with reduced neuroticism and agreeableness. This research provides insights into the psychology of elite soccer players, holding significance for talent identification, development strategies in soccer and offering insights into the psychological profiles associated with success.

**Significance statement:** This study explores the psychological profiles of elite soccer players, revealing that success on the field goes beyond physical ability. By analysing a sample of 328 participants, including 204 elite soccer players from the top teams in Brazil and Sweden, we found that elite players have exceptional cognitive abilities, including improved planning, memory, and decision-making skills. They also possess personality traits like high conscientiousness and openness, along with reduced neuroticism. Using artificial intelligence, we identified unique psychological patterns that could help in talent identification and development. These insights can be used to better understand the mental attributes that contribute to success in soccer and other high-performance fields.

## Introduction

Soccer is arguably the most widely followed sport worldwide ^1,2^, and many dream of becoming soccer players. However, only a few manage to achieve this dream. This phenomenon has cast a huge spotlight on elite soccer players, who possess exceptional skills to rise above the rest and succeed.

Originally, such attention was focused on their exceptional physical abilities ^3^. For instance, the importance of soccer players’ aerobic endurance and strength, including speed and power, has been well-documented ^4^. Similarly, it has been shown that elite female soccer players possess above-average aerobic capacities and are generally taller and heavier compared to the average female population ^5^.

Over the past decade, a new perspective has emerged, suggesting that being an elite soccer player may not solely depend on athletic capacities. Instead, it may be crucial to have a deep understanding of the game, process information quickly, and make decisions faster than others ^6^. In other words, psychological aspects ^7–9^ have become essential to comprehensively understanding elite soccer players.

Initially, researchers attempted to study sport-specific cognitive abilities ^10–14^. However, these strategies have not elucidated how specific cognitive core capacities differ from those of the general population. As a development, executive functions, i.e. higher order top-down regulatory mechanisms controlling low level processes ^15,16^, were hypothesised to be of great importance for success in ball sports as they make it possible to adapt and plan behaviour in a quickly changing environment ^17^. In line with this idea, several studies have collectively suggested that soccer players exhibit higher executive functions compared to the general population. In pioneering research, Vestberg and colleagues ^17^ found that the overall capacity for executive functions, particularly cognitive flexibility, was higher in a few elite soccer players than in sub-elite players and the general population. Notably, the primary measure of cognitive flexibility, design fluency, predicted the number of goals and assists over the following 2.5 years. Subsequent studies have reported similar findings ^18–25^, although they mostly focused only on junior players ^18,20,23,25^ or non-elite players not competing in the highest leagues ^20–22,24^ and used tests lacking a universal norm. In a preliminary attempt to solve these limitations, Vestberg and colleagues ^24^ showed that the design fluency performance was higher among national team players (NTP) compared to Premier League players (PLP), with design fluency abilities correlating with season-long assist tallies. This study also showed that coach-rated game intelligence - describing a player’s ability to read the game, adapt quickly, and always be in the right place - was associated with design fluency scores. A similar relationship between executive functions and coach-rated game intelligence was suggested in another thorough study, which however focused on a youth academy’s talent development program and included only a few professional elite players ^25^. Thus, taken together, while these studies provided valuable results, they were limited by their restricted samples of truly elite soccer players.

Furthermore, while executive functions are only part of the comprehensive psychological profile of individuals, only a couple of studies have simultaneously investigated additional psychological features in soccer players ^26,27^. For example, Vaughan and Edwards ^26^ focused on both executive functions and personality traits, revealing that soccer players exhibited heightened levels of Extraversion, Openness, and Conscientiousness, alongside superior executive function scores. Conversely, non-athletes reported elevated levels of Neuroticism and Agreeableness across all categories.

In conclusion, while previous studies have provided relevant insights, they have mostly relied on small or non-elite samples and should therefore be considered preliminary in the context of cognition in elite soccer players. To date, no large-scale study on adult elite soccer players has investigated executive functions and their relationship to successful play. Even fewer studies, if any, have extended this investigation to other psychological traits. This limitation arises from the difficulty of collecting data from truly elite soccer players, who are largely inaccessible to researchers and the general public. Additionally, previous studies did not utilise the advanced machine learning techniques available today. Consequently, a comprehensive description of the psychological profile of elite soccer players has not yet been achieved.

In this study, we addressed these gaps by taking three key steps.

First, we recruited a large sample of elite soccer players. Specifically, out of the 328 participants described in this study, 204 were elite soccer players from the top teams in the first divisions of Brazil and Sweden. Notably, this sample also included elite female players, a group often previously neglected, thereby partially addressing the usual gender bias that undermines women in football.

Second, recognising that soccer is a complex sport arguably requiring a broad spectrum of cognitive abilities and personality traits ^6,28^, and not just the executive functions predominantly investigated in previous studies ^17,18,21,22,24,29^, we expanded our investigation. We selected essential tests that effectively exemplify various aspects of the individual’s psychological profile, including planning and problem-solving, memory, executive functions, and personality traits.

Third, we complemented traditional analyses which independently assess each psychological measure with artificial neural networks. This approach provides a multivariate, comprehensive understanding of the intricate connections between our measured psychological features and constructs a model of the ideal psychological profile of elite soccer players.

In brief, our results showed that elite soccer players demonstrate heightened planning, problem-solving and memory capacities, alongside enhanced executive functions, especially cognitive flexibility. They also exhibit elevated levels of Conscientiousness, Extraversion, and Openness to experience, coupled with reduced Neuroticism and Agreeableness. Furthermore, the artificial neural network we implemented leveraged the connections between these features to build a model of the ideal psychological profile of an elite soccer player. This model successfully classified elite soccer players from matched controls with nearly 97% accuracy.

## Results

### Overview of the study

In this study, we recruited 204 elite soccer players and 124 controls matched for social and educational background. To achieve a comprehensive yet parsimonious description of the elite soccer player’s psychological profile, we selected an array of state-of-the-art psychological tests and questionnaires. These included measures for personality traits (Big Five Inventory), planning and problem-solving abilities (accuracy and speed measures of the Tower of Hanoi), working memory skills (Wechsler Adult Intelligence Scale – IV, forward and backward scales), and executive functions (Trail Making Test, Design Fluency, Colour-Word Interference, 5-Point Test). Our analyses comprised two complementary methods (as graphically depicted in **Figure 1**). First, we conducted independent analyses for each psychological feature considered in the study (i.e., personality, executive functions, planning and problem solving, and memory measures), contrasting the elite soccer players with the controls (or normative values of the tests, **Figure 2**). Second, we performed multivariate analysis using machine learning to derive the ideal, comprehensive psychological profile of the elite soccer player (**Figure 3**). Finally, by combining the measured psychological features, we predicted soccer-oriented achievements such as the number of goals, assists, and successful dribbles based on the psychological profile of individual soccer players.

**Figure 1.**
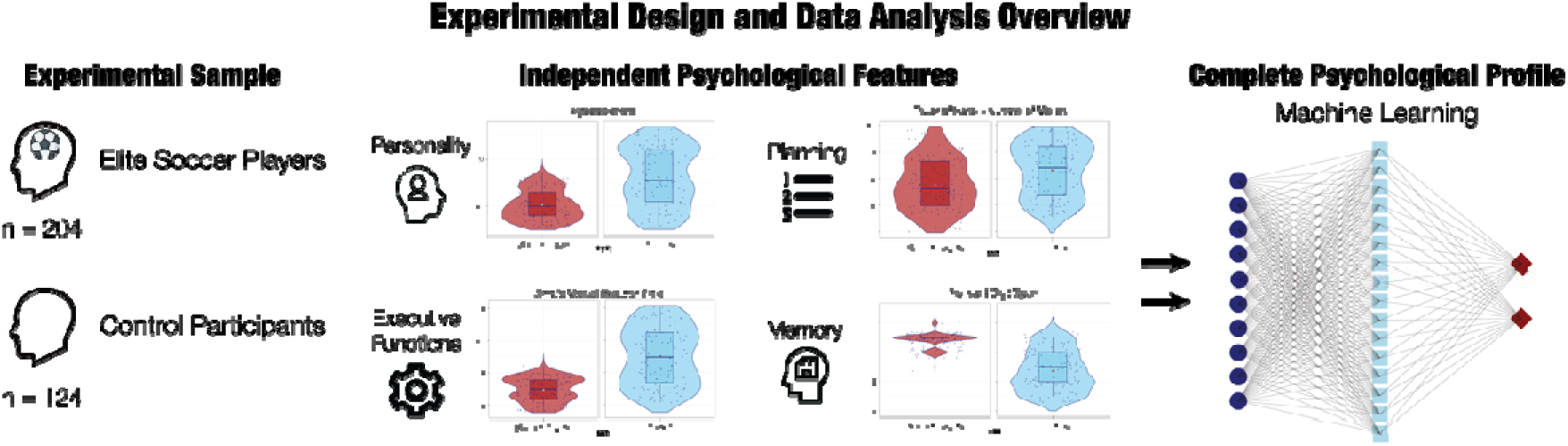
Data analysis overview. The experimental sample consisted in 204 elite soccer players and 124 controls. First, we computed independent analyses for each of the psychological features considered in the study (i.e. personality, executive functions, planning and memory measures), contrasting the elite soccer players versus the controls (or normative values of the tests). Second, we computed multivariate analysis by employing machine learning to obtain the ideal, complete psychological profile of the elite soccer player.

**Figure 2.**
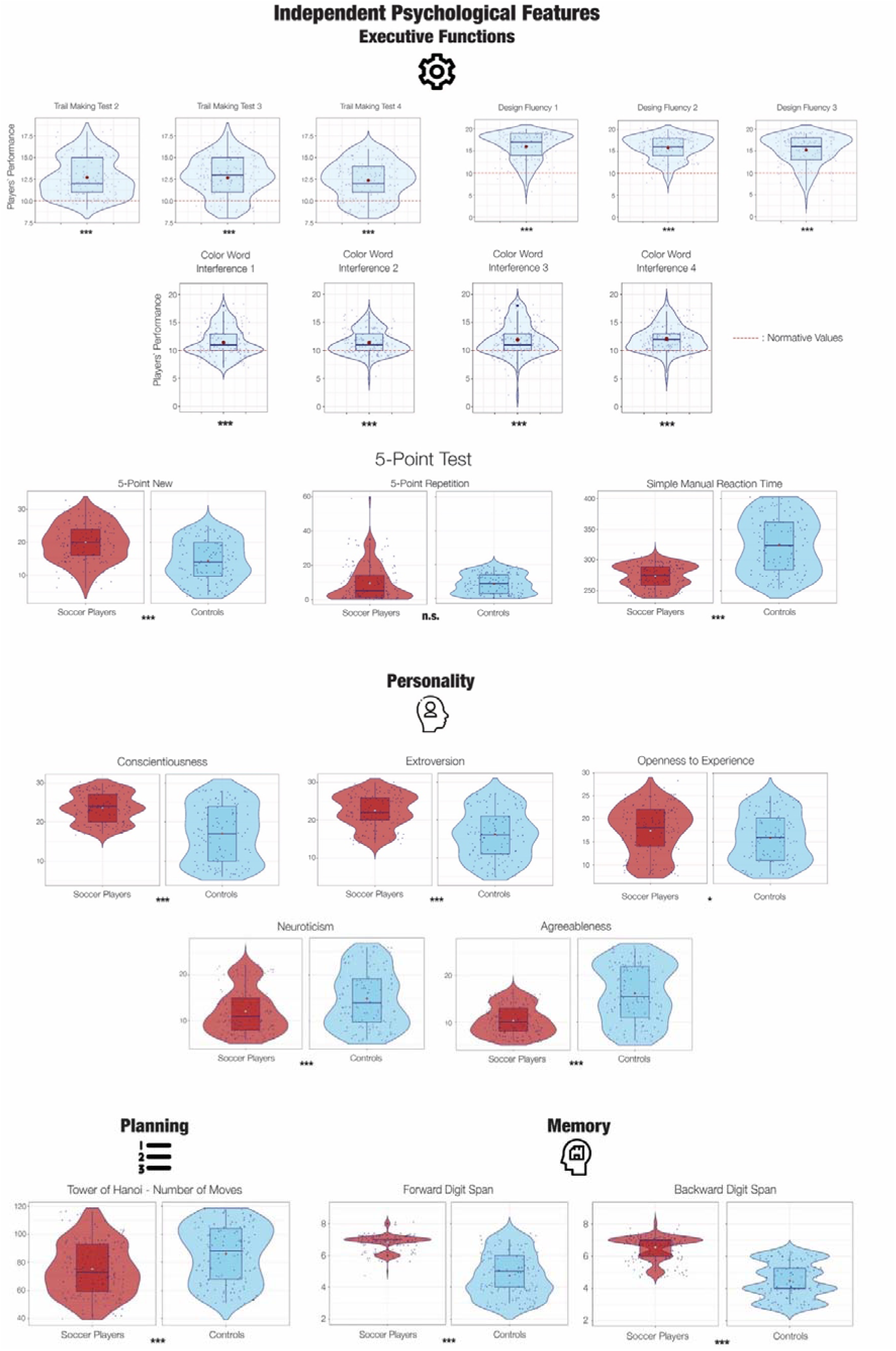
Executive functions, personality, planning and problem-solving, and memory abilities of elite soccer players. In the top part of the figure the performance of soccer players (both Brazilian and Swedish, n = 174) compared to normative values of the standard age-matched population are shown along three tests indexing executive functions: Trail Making Test, Design Fluency and Color Word Interference. The normative values are shown by red dash lines. The soccer players performance is displayed using violin plots, boxplots and scatter plots (each blue dot represents a participant). The mean across the soccer players is indicated by a larger red dot. The difference across the performance was tested using two-sample t-tests and Bonferroni corrected for multiple comparisons (p-values: * < .05; ** < .01; *** < .001; n.s. = not significant). In the bottom part of the figure, we show planning and problem-solving, memory abilities and personality scores of elite soccer players and controls matched for age and socio-economic status using several tests: 5-Point Test, Tower of Hanoi, Working Memory Test and the five dimensions of the Big Five Personality Inventory: Conscientiousness, Extroversion, Openness to Experience, Neuroticism and Agreeableness. The cognitive scores for the different tests are displayed using violin plots (red for soccer players and blue for controls), boxplots and scatter plots (each dark blue dot represents a participant). The mean across the elite soccer players is indicated by a larger light blue dot, while the mean for the controls by a red dot. The difference across the cognitive score was tested using Wilcoxon tests and Bonferroni corrected for multiple comparisons (.05 divided by five); p-values: * < .05; ** < .01; *** < .001; n.s. = not significant).

**Figure 3.**
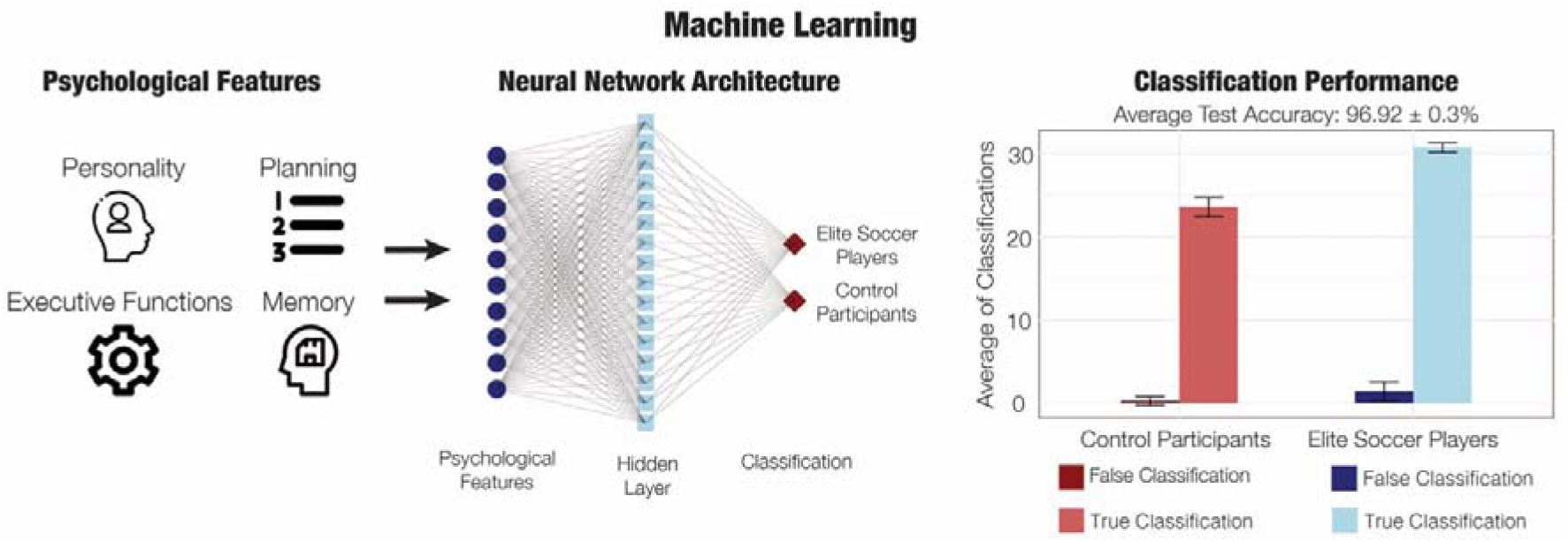
The psychological profile of the elite soccer player. The personality, planning and problem-solving, executive functions and memory scores of the elite soccer players and controls have been used to train and test a cross-validated (1000 permutations) artificial neural network algorithm aimed at classifying the elite players and the controls based solely on their combined psychological features. The classification performance of the algorithm reached almost an average 97% of accuracy in the 1000 permutations, highlighting the ideal psychological profile of the elite soccer player.

### Cognitive and personality differences between soccer players and controls

To assess the performance of soccer players in Trail Making Test, Design Fluency Test and Color-Word Interference Test from the Delis–Kaplan Executive Function System (D-KEFS) test battery in comparison to standardised norms, we computed ten two-sample t-tests and corrected for multiple comparisons using Bonferroni correction. In this analysis, 123 Brazilian and 51 Swedish elite soccer players were included who had performed the tests. The results were highly significant, as reported in **Table 1**, highlighting that soccer players were characterised by better cognitive scores than controls. We also performed independent analysis of the Brazilian (n = 123) and Swedish (n = 51) cohorts of elite soccer players to control that the results were not driven by one specific cohort (**Table S1**).

**Table 1.** Enhanced executive functions in soccer players. Results for the ten independent two-sample t-test used to assess whether the performance of soccer players in Trail Making Test, Design Fluency Test and Stroop Test was different in comparison to standardised norms, indexing the score of the average population. The table shows the tests (and the relative subscales), degrees of freedom, t-value, p-value 95% low and high confidence intervals, mean and norm.

| Test | Df | t-value | p-value | 95% CI low | 95% CI high | Mean | Norm |
| --- | --- | --- | --- | --- | --- | --- | --- |
| *TMT-1* | 173 | 15.83 | $< 2.2*10^{-16}$ | 12.37 | 13.05 | 12.71 | 10 |
| *TMT-2* | 173 | 14.07 | $< 2.2*10^{-16}$ | 12.29 | 13.03 | 12.66 | 10 |
| *TMT-3* | 173 | 14.36 | $< 2.2*10^{-16}$ | 12.04 | 12.69 | 12.36 | 10 |
| *DF-1* | 173 | 25.15 | $< 2.2*10^{-16}$ | 15.49 | 16.42 | 15.95 | 10 |
| *DF-2* | 173 | 29.16 | $< 2.2*10^{-16}$ | 15.47 | 16.26 | 15.86 | 10 |
| *DF-3* | 173 | 20.74 | $< 2.2*10^{-16}$ | 14.76 | 15.76 | 15.26 | 10 |
| *CWI -1* | 173 | 7.32 | $8.74*10^{-12}$ | 11.06 | 11.84 | 11.45 | 10 |
| *CWI -2* | 173 | 8.60 | $4.71*10^{-15}$ | 11.07 | 11.70 | 11.39 | 10 |
| *CWI -3* | 173 | 9.11 | $< 2.2*10^{-16}$ | 11.51 | 12.35 | 11.93 | 10 |
| *CWI -4* | 173 | 11.68 | $< 2.2*10^{-16}$ | 11.72 | 12.42 | 12.07 | 10 |

To assess the differences between soccer players and matched controls on cognitive and personality scores, a series of MANCOVA and Wilcoxon tests were computed, and Bonferroni corrected for multiple comparisons (see Methods for details). The Brazilian cohorts were included in this analysis (153 elite soccer players and 124 controls) as the Swedish cohort only contained elite soccer players and only performed D-KEFS tests.

There was a statistically significant difference between the elite players and controls with regards to personality scores (*F*(1, 274) = 65.70, *p* = < 2.2e-16, partial η^2^ = .55). Follow-up ANCOVAs showed statistically significant differences between the participants for all personality variables. These analyses indicated that soccer players had higher scores in Extraversion (*F*(1, 274) = 96.12, *p* < 2.2e-16, partial η^2^ = .26; mean soccer players: 22.49 ± 4.38; mean controls: 16.14 ± 6.42), Openness to experience (*F*(1, 274) = 5.29, *p* = .02, partial η^2^ = .02; mean soccer players: 17.39 ± 5.27; mean controls: 15.90 ± 5.29), Conscientiousness (*F*(1, 274) = 97.15, *p* < 2.2e-16, partial η^2^ = .26; mean soccer players: 23.49 ± 3.70; mean controls: 17.01 ± 7.17) compared to controls. On the contrary, controls reported higher scores in Neuroticism (*F*(1, 274) = 17.76, *p* = 3.4e-05, partial η^2^ = .06; mean soccer players: 12.12 ± 4.74; mean controls: 14.89 ± 6.14) and Agreeableness (*F*(1, 274) = 100.05, *p* < 2.2e-16, partial η^2^ = .27; mean soccer players: 10.28 ± 3.11; mean controls: 16.18 ± 6.26) compared to soccer players.

In relation to the Five Point Test, we computed three independent Wilcoxon tests and Bonferroni corrected them for multiple comparisons. This allowed us to assess the difference between soccer players and controls across the scores recorded for each of the variables obtained from the Five Point Test: 5-point new, 5-point repetition and simple manual reaction time. The test for 5-point new was significant (W = 13885, *p* = 3.08e-11, effect size: 0.399, soccer players sample size: 153; mean: 20.03 ± 5.92, controls sample size: 124; mean: 14.47 ± 6.15). The test for 5-point repetition was not significant (*p* = 0.17). The test for the simple manual reaction time was significant: W = 3280, *p* < 2.2e-16, effect size: 0.563, soccer players sample size: 153; mean: 273.60 ± 18.37, controls sample size: 124; mean: 324.60 ± 44.74.

With regards to WM, we computed two independent Wilcoxon tests and Bonferroni corrected them for multiple comparisons. The test for WM forward was significant: W = 16990, *p* < 2.2e-16, effect size: 0.722, soccer players sample size: 153; mean: 6.82 ± .51, controls sample size: 124; mean: 4.73 ± 1.36. The test for WM backward was also significant: W = 17594, *p* < 2.2e-16, effect size: 0.759, soccer players sample size: 153; mean: 6.54 ± .73, controls sample size: 124; mean: 4.47 ± 1.11.

In relation to the Tower of Hanoi Test, we computed two independent Wilcoxon tests (one for the seconds and one for the moved used to complete the task) and Bonferroni corrected them for multiple comparisons. The test for the number of seconds used to complete the task was not significant (*p* = .33). The test regarding the number of moves was significant, indicating that soccer players needed a lower number of moves to complete the task: W = 6630.5, *p* = 1.65e-05, effect size: 0.259, soccer players sample size: 153; mean: 75.19 ± 18.90, controls sample size: 124; mean: 86.08 ± 20.29.

### Cognitive and personality profile of soccer players and soccer performance

To investigate the relationship between personality traits, cognitive scores, and a selected array of specific soccer skills/objectives, five general linear model (GLM) analyses were conducted, and Bonferroni corrected for multiple comparisons. The independent variables for the five GLMs included the cognitive and personality scores which were significant in the previous tests, while the dependent variables, inserted separately in the five GLMs, were: the number of goals scored by the players (i), number of shots tempted per game (ii), number of assists per game (iii) and average attempted (iv) and successful dribbles (v). The Brazilian cohort (n = 153) was included in this analysis since this data was not available for the Swedish cohort. The results are reported in **Table 2**.

**Table 2.** Psychological features predict soccer performance. Five general linear models (GLMs) assessing the predictive power of personality and cognitive scores on a selected array of soccer skills/objectives. The table reports the key statistics for each of the five models, as well as the p-values and coefficients (only if significant) for independent variables (predictors).

|  | Dependent variables |  |  |  |  |
| --- | --- | --- | --- | --- | --- |
|  | GLM 1 - Goal | GLM 2 - shot | GLM 3 - assist | GLM 4 - att. dribb. | GLM 5 - succ. dribb. |
| **Model statistics** | Adj. R<sup>2</sup> = .22<br>*p* = 1.22e-06 | Adj. R<sup>2</sup> = .07<br>*p* = .03 | Adj. R<sup>2</sup> = .08<br>*p* = .02 | Adj. R<sup>2</sup> = .29<br>*p* = 3.25e-09 | Adj. R<sup>2</sup> = .31<br>*p* = 7.04e-10 |
| **Predictors** |  |  |  |  |  |
| *Age* | *p* = .33 | *p* = .15 | *p* = .84 | *p* = .28 | *p* = .11 |
| *Neuroticism* | *p* = .25 | *p* = .07 | *p* = .06 | *p* = .02<br>$\beta$ = -.07 | ***p* = .009**<br>**$\beta$ = -.05** |
| *Extraversion* | *p* = .60 | *p* = .12 | *p* = .30 | *p* = .50 | *p* = .65 |
| *Openness* | ***p* = .003**<br>**$\beta$ = .16** | *p* = .27 | ***p* = .008**<br>**$\beta$ = .08** | ***p* = 2.03e-06**<br>**$\beta$ = .17** | ***p* = 7.71e-07**<br>**$\beta$ = .09** |
| *Agreeabl.* | *p* = .48 | *p* = .35 | *p* = .60 | *p* = .67 | *p* = .96 |
| *Conscient.* | ***p* = .04**<br>**$\beta$ = -.15** | *p* = .18 | *p* = .49 | *p* = .17 | *p* = .43 |
| *5-point new* | ***p* = 6.8e-07**<br>**$\beta$ = .24** | ***p* = .01**<br>**$\beta$ = .03** | *p* = .21 | *p* = .42 | *p* = .67 |
| *S Manual RT* | *p* = .44 | *p* = .71 | *p* = .68 | *p* = .84 | *p* = .78 |
| *WM* | *p* = .28 | *p* = .69 | *p* = .39 | ***p* = .002**<br>**$\beta$ = 6.94** | ***p* = .005**<br>**$\beta$ = 3.69** |
| *Hanoi* | *p* = .23 | *p* = .74 | *p* = .76 | ***p* = .02**<br>**$\beta$ = .02** | ***p* = .0004**<br>**$\beta$ = .02** |

### Artificial neural network performance

The neural network model, trained and evaluated using permutation-based (1000 permutations) cross-validation, demonstrated robust predictive capabilities for assessing the relationship between personality traits, cognitive scores, and soccer membership. Average Test Loss: 0.14 ± .04; Average Test Accuracy: 96.92 ± .03 %; Average Precision: 95.66 ± .03 %; Average Recall: 99.06 ± .02 %; Average F1 Score: 97.29 ± .02 %. The average confusion matrix, summarising the model’s classification performance, is presented as follows and illustrated in **Figure 3**. It includes the distribution of true positive (TP): 30.71 ± .55; true negative (TN): 23.57 ± 1.14; false positive (FP): 1.43 ± 1.14; and false negative (FN): .29 ± .55 predictions, providing a detailed insight into the model’s performance across different classes. The model achieved a high level of precision, recall, and F1 score, indicative of its ability to effectively distinguish between soccer players and control subjects. These results suggest that this neural network model reliably predicted professional soccer membership based on personality and cognitive factors. This analysis involved the Brazilian cohorts (153 elite soccer players and 124 controls) since it required participants with no missing scores across the personality and cognitive tests.

## Discussion

Our investigation unveiled notable differences in the performance of elite soccer players versus matched controls and the norm across several cognitive tests and personality ratings. This allowed us to delineate the psychological profile of elite soccer players. These athletes displayed superior cognitive abilities, including higher levels of executive functions, memory, problem-solving, and planning. Additionally, their personality profile diverged with elevated scores in Extraversion, Openness to Experience, and Conscientiousness, alongside lower scores in Agreeableness and Neuroticism. The tests were also able to predict several successful soccer behaviours ranging from goals and assists to different types of dribbling. Finally, the artificial neural network we implemented leveraged the connections between the measured psychological features to build a model of the ideal psychological profile of an elite soccer player, successfully classifying elite soccer players from matched controls with nearly 97% accuracy.

The psychological profile that emerged from our results of heightened executive functions in elite soccer players aligns with prior studies ^17,18,21,22,24^. Our findings expand on this by highlighting superior performance in cognitive flexibility and inhibition response within an extensively larger sample of elite soccer players compared to previous studies. Notably, while some earlier studies included a relatively large number of participants ^21–23,25^, most of them were not adult nor elite players competing in a premiere league. Therefore, while previous results are valuable yet arguably preliminary, the present findings are more conclusive. Interestingly, similar results have also been reported in other team sports, including volleyball ^30^, ice hockey ^31^, and basketball ^32^. Despite variations in the role of executive functions across different sports ^33^, these studies collectively suggest a consistent trend: elite athletes generally outperform their less experienced counterparts in executive functions ^34–37^.

Remarkably, previous studies have primarily focused on a limited set of executive functions, yet soccer proficiency encompasses a broader spectrum of abilities. This includes memory, visuo-spatial reasoning, and planning ^6,28^. In our study, we demonstrate that elite soccer players significantly outperform matched controls in memory, problem-solving, and planning tasks. Previous research on these psychological traits within soccer is rather sparse but suggests that soccer players might be characterised by higher levels of working memory compared to controls ^18,22^. Our findings are consistent with and expand on these studies, underscoring the pivotal role of memory processes in soccer performance in a large sample of elite soccer players. Furthermore, the current research investigated strategic planning using the Tower of Hanoi as an essential assessment tool to obtain insights into participants’ spatial reasoning and problem-solving abilities. Our results indicate that soccer players clearly outperformed controls, suggesting superior efficiency in analysing situations and devising optimal strategies to achieve their goals in the same time frame. The ability to plan several steps ahead in order to reach a goal in a quickly changing environment may be one of the most crucial cognitive processes related to successful behaviour in complex ball sports such as soccer.

In our study, we assessed personality traits and found that elite soccer players exhibit heightened levels of imagination, open-mindedness, self-discipline, and energy. These findings are consistent with prior research on personality traits in elite athletes in general ^38^ and a systematic review examining the relationship between personality and athletic performance ^39^. Our study expands on this body of research by specifically focusing on elite soccer players. Additionally, our study revealed lower levels of cooperativeness and a reduced propensity for negative emotions, as evidenced by lower scores in Neuroticism and Agreeableness. This contrasts with findings from previous studies on other sports, suggesting that soccer players’ personality profiles differ from those of different categories of athletes. Specifically, soccer players tend to score particularly low in Agreeableness. This trend may be attributed to the global popularity of soccer, which fosters intense competition at elite levels where being particularly agreeable could represent a disadvantage. In broader research on personality traits among athletes, other studies have employed different yet compatible measurement tools. For example, Gabrys and Wontorczyk ^40^ used temperamental and impulsivity scales in their study of short-track speed skating to achieve a more nuanced understanding of athlete profiles. They reveal that these athletes are characterised by high scores on temperamental scales such as persistence, harm avoidance, and novelty seeking, as well as character scales like cooperativeness. Additionally, they score high on impulsivity scales, including sensation seeking and positive urgency. Similarly, Kang et al. ^41^ explored temperamental factors to elucidate success among baseball players, suggesting that predictive temperamental factors for success in these athletes include traits such as novelty seeking and reward dependence.

An innovative aspect of our study is the application of artificial neural networks, which shows promise for future research. We applied this approach to differentiate between elite soccer players and controls based solely on cognitive and personality attributes. Our findings demonstrate the efficacy of this method in accurately predicting participants’ membership in these groups. This suggests that the psychological profile of soccer players exhibits distinctive characteristics effectively captured by our model. Specifically, the confusion matrix of our classifier reveals near-perfect accuracy in predicting soccer players based on their psychological features. Among the few errors identified by our model, we observed that almost all were false positives, where individuals who were not athletes were incorrectly classified as athletes. This finding is particularly intriguing and logical, suggesting that successful soccer players possess a distinct psychological profile. Essentially, without this specific psychological profile, achieving success in soccer appears extremely improbable. Conversely, possessing the correct psychological profile does not guarantee success, as various factors such as personal choices, preferences, or physical limitations can still hinder one’s soccer career.

A final interesting result delivered by our research lies in identifying specific abilities that serve as predictors of future performance outcomes. Our findings build upon previous studies by Vestberg and colleagues ^17,18,24^, which demonstrated significant correlations between working memory and goals scored ^18^, design fluency capacity and assists made ^24^, and executive function and goals/assists scored in subsequent seasons ^17^. In our study, utilising different predictors and performance indicators, we discovered additional significant relationships. Importantly, we considered both cognitive and personality factors as predictors and examined a diverse range of soccer abilities beyond goals and assists, such as successful dribbles. Future research is needed to further explore these aspects, as such insights could enhance sports practices and assist coaches in identifying individuals likely to achieve optimal performance.

Beyond elite soccer players and athletes in general, the findings of our study can also be viewed in the context of broader human success and psychology. While we have described the essential psychological traits for becoming an elite soccer player, an intriguing question arises: how do these results apply to other domains and predict success in those areas? Interestingly, our findings resonate beyond sports. For instance, research focusing on chess ^42^ has shown that expert players demonstrate significantly higher intelligence compared to non-players, correlating their playing proficiency with cognitive abilities. This parallels our observations, where elite soccer players competing at the highest levels exhibit exceptional cognitive abilities. Moreover, studies on professional gamers have similarly reported superior performance in cognitive assessments ^43,44^ and enhanced cognitive flexibility ^45^, aligning with our findings in elite soccer athletes. These cognitive advantages are also evident in other domains of expertise; professional musicians, for instance, outperform controls in cognitive assessments ^46–49^. This suggests that expertise in music, similar to elite sports, is associated with heightened cognitive abilities. Personality traits have also been studied across various domains beyond sports, remaining crucial for achieving success in professional contexts. For example, our findings align with studies on professional politicians, which reported elevated levels of Energy and Agreeableness. Nettle ^50^ explored personality traits in professional actors compared to the general population, reporting a profile which is similar to the one described in the current study soccer players, particularly in relation to higher levels of Extraversion and Openness to Experience. However, contrary to our results, Nettle found that professional actors also exhibited higher levels of Agreeableness, challenging the notion that top-level performance requires lower Agreeableness levels. Taken together, our results provide insights not only into elite soccer players but also contribute, alongside findings from previous literature, to a broader psychological profile associated with achieving success in the society.

In conclusion, this study delineated the psychological profile of elite soccer players through comprehensive analyses of cognitive and personality measures in a large sample. Our findings not only define this profile but also shed light on the broader psychological traits which might be crucial for success in society. Looking towards future perspectives, our results reignite the enduring debate of nature versus nurture. Are the psychological traits identified in this study innate or acquired? Furthermore, can deliberate interventions during their developmental stages shape the optimal psychological profile of elite soccer players? Addressing these questions through carefully designed longitudinal experiments represents a crucial objective for future research, building upon the current study to deepen our understanding of human behaviour and provide insights into fostering excellence in professional soccer and other elite human activities.

## Methods

### Participants

The sample consisted of three groups of participants.

The first cohort was composed of 153 Brazilian elite soccer players (0 females, 153 males, mean age: 26.14 ± 5.63 years, average years of formal education: 8.74 ± 1.83) who played in the Brazilian major league at the time of the data collection. The second cohort was composed of 124 Brazilian control participants (43 females, 34 males, mean age: 27.38 ± 5.25 years, average years of formal education: 8.01 ± 1.82). The third cohort was composed of 51 professional Swedish soccer players (19 females, 32 males, mean age: 24.50 ± 4.60 years, age range 17 to 35 years; Men: 24.50 ± 4.70 years, age range 18 to 35 years; Females: 24.50 ± 4.60, age range 17 to 33 years). Within this group of 51 players, 28 individuals had not previously participated in any senior-level matches for a national team, designated as Premier League Players (PLP). Meanwhile, the remaining 23 players, classified as National Team Players (NTP), had experience in playing at least one game for their respective senior national teams. Notably, these 23 NTPs had represented a total of 14 different national teams globally. While preliminary results from this third cohort have been previously presented ^24^, in the current study they were gathered with the first two larger cohorts to obtain an extensive global sample of adult elite soccer players, which allowed us to draw more solid and definitive conclusions. As a matter of fact, for certain analyses, a combined dataset of Swedish and Brazilian players was created and juxtaposed against norm values derived from general populations. In other analyses, the Brazilian elite soccer players were specifically contrasted with the control group. The project was approved by the following ethical committees: Regionala etikprövningsnämn - Stockholm; Dnr 2017/2453-31/5 (Swedish cohort) and Rio de Janeiro State University Ethical Committee under the consubstantiated report 2.990.037 (Brazilian cohorts). The experimental procedures complied with the Declaration of Helsinki – Ethical Principles for Medical Research. Participants’ informed consent was obtained before starting the experiment.

### Psychological tests and procedure

In our study, we employed multiple assessments to evaluate the personality and cognitive scores of the participants. This approach enabled us to derive a comprehensive understanding of the cognitive and personality profiles of professional soccer players in comparison to controls who were matched for age and education.

The study combined three international datasets in which participants were tested either in Sweden or in Brazil. Regardless of the place, the assessment of each participants occurred independently within a quiet and controlled environment. The study provided a dedicated room equipped with the necessary resources, where participants could complete the several tests we administered without external disturbances. An experimenter was present and readily available throughout the assessment sessions to address any queries or concerns that participants might have encountered. Participants were given standardised instructions and encouraged to respond honestly and to the best of their possibilities to each item of the tests. As follows, we provide detailed information on the tests used, specifying which samples of participants underwent specific assessments.

The Big Five Inventory (BFI) ^51^ questionnaire was used to assess participants’ personality traits across five dimensions: Openness to Experience, Conscientiousness, Extraversion, Agreeableness, and Neuroticism. The BFI is a widely employed self-report instrument that has demonstrated good reliability and validity in previous studies and has been widely used across various cultures and populations. The BFI was administered to the Brazilian cohorts (153 elite soccer players and 124 controls).

For the assessment of participants’ cognitive abilities, we utilised a battery of tests aimed at constructing a thorough profile, particularly focusing on executive functions, visuo-spatial reasoning, memory, inhibition control, and the ability to follow traces. This comprehensive approach allowed us to investigate the nuanced aspects of cognitive functioning among soccer players, shedding light on specific domains that contribute to their cognitive profiles.

We employed three scales from the Delis-Kaplan Executive Function System (D-KEFS) ^52^: the Design Fluency Score, Colour-Word Interference, and Trail Making Test. The D-KEFS is a comprehensive neuropsychological assessment tool designed to evaluate various aspects of executive functioning, such as cognitive flexibility, inhibition control, and problem-solving. The Design Fluency Score assesses participants’ capacity for generating as many novel designs as possible within a set period, providing insights into creative thinking and mental flexibility. In our study, this score comprised three progressively challenging subscales. In the initial stage, participants were tasked with connecting provided black dots exclusively into novel four-line shapes (the test is mainly focusing on cognitive flexibility but also includes behavioural flow, convergent creativity, response inhibition and working memory). Subsequently, the difficulty increased as participants were required to connect white dots, disregarding the presence of black dots (this variant add a need for behavioural inhibition). The third and final stage introduced a more complex task, where participants alternated between connecting one black and one white dot in a sequential manner (this variant increases the need for cognitive flexibility). This nuanced progression in difficulty allowed us to capture a spectrum of cognitive abilities related to design fluency, ranging from basic pattern recognition to more complex cognitive flexibility. The Colour-Word Interference test is a Stroop-task and serves as a robust measure of inhibition control and cognitive flexibility. In our study, this test was administered through four consecutive short blocks, each presenting participants with tasks of a similar nature. These tasks required participants to inhibit automatic responses to conflicting stimuli (e.g. participants were requested to name the colour of a colour-word which was printed in a different, conflicting colour). The task offers an evaluation of their ability to adapt and flexibly switch between cognitive processes. Finally, the Trail Making Test was employed to gauge visual attention, sequencing, and motor speed. This test featured three distinct subscales: first, connecting numbers in ascending order; second, linking letters in alphabetical order; and third, alternating between connecting one number and one letter in sequence. The inclusion of these subscales allowed for a comprehensive evaluation of visual attention and cognitive processing speed, providing valuable insights into participants’ abilities to navigate sequential tasks with varying complexity. The selection of these specific D-KEFS scales was deliberate, as they collectively offer a nuanced perspective on the executive functions essential for high-performance activities like professional soccer. These tests were administered to both the Brazilian (n = 123 elite soccer players out of the total n = 153 since 30 players did not complete the D-KEFS battery) and Swedish (n = 51 elite soccer players) cohorts.

The Five-Point Test ^53^ is a structured and standardised assessment designed to measure figural fluency functions. In our study, we utilised three distinct measures to estimate participants’ performance on this test: the 5-Point New, 5-Point Repetition, and Simple Manual Reaction Time. The 5-Point New involves participants generating unique designs in response to a set of specific criteria within a five-point framework. This measure assesses participants’ ability to produce novel and varied figural patterns, offering insights into creative thinking and cognitive flexibility. On the other hand, the 5-Point Repetition requires participants to replicate a given design within the same five-point structure, providing a measure of precision and attention to detail. Additionally, the Simple Manual Reaction Time measure evaluates participants’ basic motor speed and reaction time during the 5-Point task by recording their response times to simple stimuli. Together, these three measures contribute to a comprehensive assessment of figural fluency functions, offering a nuanced understanding of participants’ cognitive performance in this domain. The Brazilian cohorts performed this test (153 elite soccer players and 124 controls).

Finally, we incorporated two additional cognitive assessments to comprehensively evaluate participants’ memory and planning abilities: the Forward and Backward Working Memory Tests from the Wechsler Adult Intelligence Scale-IV ^54^ and the Tower of Hanoi ^55^ task. The Forward and Backward Working Memory Tests are classic measures designed to assess the capacity to temporarily store and manipulate information in participants’ mind. The Forward version involves participants recalling and reproducing a sequence of presented numbers in the same order, while the Backward version requires them to recall the sequence in reverse order, tapping into the cognitive process of mental manipulation. The Brazilian cohorts performed this test (153 elite soccer players and 124 controls).

Furthermore, the Tower of Hanoi task was utilised, employing both the time taken to complete the task and the number of moves required as performance metrics. This task is a well-established measure of executive functions, specifically problem-solving, planning, and spatial reasoning. Participants were tasked with moving a series of disks from one peg to another, adhering to specific rules. The duration and efficiency of task completion provide valuable insights into participants’ cognitive processes related to strategic planning and problem-solving skills. The Brazilian cohorts performed this test (153 elite soccer players and 124 controls).

The comprehensive battery of cognitive assessments employed in our study was meticulously curated to yield a nuanced and thorough understanding of various facets of human cognition. By incorporating a diverse range of tests, including measures of executive functions, figural fluency, working memory, and problem-solving, we aimed to capture the multifaceted nature of cognitive abilities in professional soccer players. The inclusion of well-established and standardised tests ensures the reliability and validity of our findings, providing a robust foundation for analysing cognitive differences among participants.

### Soccer players against controls

This study employed comparative analyses to investigate whether soccer players were characterised by a different cognitive and personality profile compared to controls. As described below, this was done either by comparing the soccer players’ scores versus standardised values (using two-sample t-tests) of the population or by contrasting the soccer players’ scores against the ones of controls matched for age and socio-economic background [using one-way multivariate analysis of covariance (MANCOVA) or Wilcoxon tests when the assumptions of normality for the MANCOVAs were not met]. Independent analyses were performed grouping families of cognitive and personality variables in the same statistical models. All the statistical analyses were computed using the R statistical software (R Core Team, 2022).

First, using a series of independent two-sample t-tests, we assessed the performance of soccer players in Trail Making Test, Design Fluency Test and Stroop Test in comparison to standardised norms, indexing the score of the average population.

Specifically, ten t-tests were employed, corresponding to three scales for the Trail Making Test, three for the Design Fluency Test, and four for the Stroop Test. To address the issue of multiple comparisons, Bonferroni correction was applied, resulting in a revised threshold p-value of .005 (obtained by dividing .05 by the total number of tests, i.e., 10). Both Brazilian (n = 123 elite soccer players out of the total n = 153 since 30 players did not complete the D-KEFS battery) and Swedish (n = 51) elite soccer players (total n = 174) were included in this analysis, as they underwent the exact same set of tests.

Second, we computed five MANCOVAs [at α = .01 after applying the Bonferroni correction (.05 divided by five)] to contrast several personality and cognitive scores of soccer players versus controls. This involved only the full Brazilian cohorts (soccer players n = 153, controls n = 124).

The five personality scores taken from the Big Five Questionnaire (Neuroticism, Extraversion, Openness to Experience, Agreeableness, Conscientiousness) were tested. Unlike the previous analysis that relied solely on norms of the general population, this investigation incorporated data not only for soccer players but also for matched controls. A one-way MANCOVA (Wilk’s Lambda [Λ], adjusted α = .01) was performed to compare the personality scores between the two groups while controlling for age. The independent variable was group (soccer players and controls), and the model included five dependent variables (Neuroticism, Extraversion, Openness to Experience, Agreeableness, Conscientiousness) along with one covariate (age). Afterwards, to determine the effects of the independent variable on each of the dependent variables inserted in the MANCOVA, five univariate analyses of covariance (ANCOVA) were computed. These were computed at α = .01 after applying the Bonferroni correction (.05 divided by the number of ANCOVAs conducted) as follow-up tests to the MANCOVA.

A second MANCOVA (Wilk’s Lambda [Λ], adjusted α = .01) was computed for the Five Point Test. In this analysis, the independent variable was group (soccer players and controls), and the dependent variables comprised three variables from the Five Point Test: 5-point new, 5-point repetition, and simple manual reaction time. Once again, age served as the covariate. Although the MANCOVA yielded statistical significance, the residuals were found to deviate from linearity. Consequently, three independent Wilcoxon tests were executed separately for each of the aforementioned variables (5-point new, 5-point repetition, and simple manual reaction time) to compare performance between soccer players and controls. To account for multiple comparisons, a Bonferroni correction was applied, resulting in an adjusted threshold *p-value* of .016 (obtained by dividing .05 by the number of independent Wilcoxon tests conducted, i.e., 3).

Subsequently, we computed a MANCOVA (Wilk’s Lambda [Λ], adjusted α = .01) for the WM test. Here, the independent variable was group (soccer players and controls), the dependent variables were forward digit span and backward digit span, and the covariate was the age of the participants. Once again, although the MANCOVA was significant, its residuals were not linearly distributed. Thus, we computed two independent Wilcoxon tests to compare, separately, each of the variables reported above (forward digit span and backward digit span) across soccer players and controls. Bonferroni correction for multiple comparisons was applied, obtaining a new thresholding *p-value* corresponding to .05/2 = .025.

A last MANCOVA (Wilk’s Lambda [Λ], adjusted α = .01) was computed for the Tower of Hanoi test. Here, the independent variable was group (soccer players and controls), the dependent variables were seconds (time) and number of moves used for the completion of the task, and the covariate was the age of the participants. As for the previous analysis, although the MANCOVA was significant, its residuals were not linearly distributed. Thus, we computed two independent Wilcoxon tests to compare, separately, each of the variables reported above (seconds and number of moves) across soccer players and controls. Bonferroni correction for multiple comparisons was applied, obtaining a new thresholding p-value corresponding to .05/2 = .025.

To be noted, Bonferroni correction was applied for adjusting the α of the five MANCOVAs described. The adjusted α corresponded to .05/5 = .01. The effect size of each MANCOVA was calculated using partial eta squared (i.e., partial η^2^), while the effect size for the Wilcoxon tests was computed using by computing Z statistic and dividing it by the square root of the sample size, as commonly done by the R statistical software (R Core Team, 2022).

### Cognitive and personality profile of soccer players and soccer performance

To investigate the relationship between personality traits, cognitive scores, and a selected array of specific soccer skills/objectives, five general linear model (GLM) analyses were conducted using the R statistical software (R Core Team, 2022). The independent variables for the five GLMs included the cognitive and personality scores which were significant in the previous tests. Specifically, the independent variables were Neuroticism, Extroversion, Openness, Agreeableness, Conscientiousness, 5-points new and simple manual reaction times, WM (forward and backward combined), and Tower of Hanoi (moves), and age were included as predictors.

We focused on five key soccer abilities/objectives, which were inserted as dependent variable, separately for the five GLMs that we computed. These skills/objectives corresponded to the number of goals scored by the players (i), number of shots attempted per game (ii), number of assists per game (iii) and average attempted (iv) and successful dribbles (v).

Since we computed five independent GLMs, we computed the Bonferroni correction for multiple comparisons, which set the significance level for hypothesis testing at *p* = .01. This analysis involved only the Brazilian cohort of elite soccer players (n = 153) since this data was not available for the Swedish cohort.

### Artificial neural networks

Artificial neural networks (ANNs) ^56^ were used to decode soccer players from controls based on their cognitive and personality profile. The implementation of the ANNs was coupled with permutation-based cross-validation to prevent overfitting and ensure robustness of the results. The dataset included the same variables described for the previous analyses and involved personality traits (Neuroticism, Extroversion, Openness, Agreeableness, Conscientiousness) and cognitive scores (5-points new, Simple Manual Reaction Time, Forward Digit Span, Backward Digit Span, Tower of Hanoi moves). The dataset was divided into soccer player and control groups, and features were normalised by scaling them between zero and one. As mentioned above, to account for potential overfitting and ensure robust model evaluation, a permutation-based cross-validation strategy was employed. The data underwent 1000 permutations, with the data randomly reassigned to either training or testing set, for each iteration. The ANN architecture was constructed using the TensorFlow and Keras frameworks in Python. The model comprised an input layer with the same number of neurons as input variables (ten), followed by a hidden layer with 16 neurons and a ReLU activation function. The output layer consisted of a single neuron with a sigmoid activation function, which is suitable for binary classification tasks. The model was trained using the Adam optimiser and binary cross-entropy loss function. During training, the data was processed in batches of 32 samples for 20 epochs, as commonly done in ANN implementations. The evaluation of the model on the test set was performed separately after training. Here, for each permutation, various performance metrics were computed and stored, including test loss, test accuracy, precision, recall, F1 score, and confusion matrix.

The test loss is a measure of how well the model is performing in predicting whether an individual is a soccer player or a control. It quantifies the disparity between the model’s predictions and the actual outcomes in the test dataset. Lower test loss values indicate a closer alignment between the predicted and actual values. Test accuracy provides a straightforward gauge of the model’s correctness in classifying instances within the test set. It represents the proportion of correctly classified samples, showcasing the model’s overall accuracy in distinguishing between soccer players and controls. Precision focuses on the accuracy of positive predictions. It assesses the ratio of correctly predicted soccer players to the total instances predicted as soccer players. A higher precision value signifies fewer instances where the model wrongly identified a control as a soccer player. Recall, also known as sensitivity or true positive rate, evaluates the model’s ability to correctly identify actual soccer players. It measures the ratio of correctly predicted soccer players to the total actual soccer players, emphasizing the model’s sensitivity in recognizing true positives. The F1 score combines precision and recall, providing a balanced metric that considers both false positives and false negatives. This harmonic mean is particularly useful when there is an imbalance in class distribution, offering a comprehensive assessment of the model’s effectiveness. The confusion matrix is a visual representation of the model’s classification performance. It delineates true positive, true negative, false positive, and false negative predictions. This matrix allows for a nuanced understanding of specific types of errors and correct classifications, aiding in the identification of areas for model improvement.

The results from the 1000 permutations were aggregated to compute average values for the aforementioned performance metrics. This analysis involved only the Brazilian cohort of elite soccer players (n = 153) since it required participants with no missing scores across the personality and cognitive tests.

## Code availability

The code used for the analyses is available in the following repository: https://github.com/leonardob92/Soccer_PsychologicalProfile.git

## Data availability

The anonymized data from the experiment will be made available upon reasonable request.

## Acknowledgements

The Center for Music in the Brain (MIB) is funded by the Danish National Research Foundation (project number DNRF117).

L.B. is supported by Lundbeck Foundation (Talent Prize 2022), Carlsberg Foundation (CF20-0239), Center for Music in the Brain, Linacre College of the University of Oxford, Society for Education and Music Psychology (SEMPRE’s 50^th^ Anniversary Awards Scheme), and Nordic Mensa Fund.

M.L.K. is supported by Center for Music in the Brain and Centre for Eudaimonia and Human Flourishing, which is funded by the Pettit and Carlsberg Foundations.

## Author contributions statement

P.P., T.V., F.A.G., R.J., L.B. and M.L.K. conceived the hypotheses. V.T., R.J., P.P. and A.F.G. designed the study. A.F.G., P.P., M.I., M.L.K. and L.B. recruited the resources for the experiment. A.F.G, R.J. and T.V. collected the data. L.B. performed statistical analysis and modelling. P.P., A.F.G., M.I. and M.L.K. provided essential help to interpret and frame the results within the scientific literature. L.B. and D.S. wrote the first draft of the manuscript. L.B. and D.S. prepared the figures. All the authors contributed to and approved the final version of the manuscript.

## Supplementary information

**Table S1.** Enhanced executive functions in soccer players – Brazilian and Swedish cohorts. Results, reported independently for the Swedish and Brazilian cohorts, for the ten independent two-sample t-test used to assess whether the performance of soccer players in Trail Making Test, Design Fluency Test and Stroop Test was different in comparison to standardised norms, indexing the score of the average population. The table shows the tests (and the relative subscales), degrees of freedom, t-value, p-value 95% low and high confidence intervals, mean and norm.

| Brazilian cohort (elite soccer players) |  |  |  |  |  |  |  |
| --- | --- | --- | --- | --- | --- | --- | --- |
| Test | Df | t-value | p-value | 95% CI low | 95% CI high | Mean | Norm |
| *TMT-1* | 122 | 12.20 | < 2.2e-16 | 12.30 | 13.19 | 12.74 | 10 |
| *TMT-2* | 122 | 12.31 | < 2.2e-16 | 12.34 | 13.24 | 12.79 | 10 |
| *TMT-3* | 122 | 12.20 | < 2.2e-16 | 12.17 | 13.00 | 12.59 | 10 |
| *DF-1* | 122 | 21.08 | < 2.2e-16 | 15.62 | 16.79 | 16.20 | 10 |
| *DF-2* | 122 | 26.23 | < 2.2e-16 | 15.94 | 16.91 | 16.42 | 10 |
| *DF-3* | 122 | 20.54 | < 2.2e-16 | 15.33 | 16.47 | 15.90 | 10 |
| *CWI-1* | 122 | 7.36 | 2.3e-11 | 11.37 | 12.37 | 11.87 | 10 |
| *CWI -2* | 122 | 7.65 | 5.3e-12 | 11.11 | 11.89 | 11.50 | 10 |
| *CWI -3* | 122 | 8.15 | 3.6e-13 | 11.61 | 12.65 | 12.13 | 10 |
| *CWI -4* | 122 | 10.63 | < 2.2e-16 | 11.87 | 12.72 | 12.29 | 10 |
| Swedish cohort (elite soccer players) |  |  |  |  |  |  |  |
| Test | Df | t-value | p-value | 95% CI low | 95% CI high | Mean | Norm |
| *TMT-1* | 50 | 12.02 | 2.3e-16 | 12.19 | 13.07 | 12.62 | 10 |
| *TMT-2* | 50 | 6.86 | 9.8e-09 | 11.66 | 13.04 | 12.35 | 10 |
| *TMT-3* | 50 | 8.43 | 3.7e-11 | 11.39 | 12.26 | 11.82 | 10 |
| *DF-1* | 50 | 14.21 | < 2.2e-16 | 14.59 | 16.11 | 15.35 | 10 |
| *DF-2* | 50 | 16.76 | < 2.2e-16 | 13.97 | 15.05 | 14.51 | 10 |
| *DF-3* | 50 | 8.19 | 8.6e-11 | 12.80 | 14.62 | 13.71 | 10 |
| *CWI-1* | 50 | 1.88 | .066 | 9.97 | 10.89 | 10.43 | 10 |
| *CWI -2* | 50 | 3.98 | .0002 | 10.54 | 11.65 | 11.10 | 10 |
| *CWI -3* | 50 | 4.14 | .0001 | 10.75 | 12.15 | 11.45 | 10 |
| *CWI -4* | 50 | 5.13 | 4.7e-06 | 10.93 | 12.13 | 11.53 | 10 |

